# High-Throughput Single-Molecule Analysis via Divisive Segmentation and Clustering

**DOI:** 10.1101/603761

**Authors:** David S. White, Marcel P. Goldschen-Ohm, Randall H. Goldsmith, Baron Chanda

## Abstract

Single-molecule approaches provide insight into the dynamics of biomolecules, yet analysis methods have not scaled with the growing size of data sets acquired in high-throughput experiments. We present a new analysis platform (DISC) that uses divisive clustering to accelerate unsupervised analysis of single-molecule trajectories by up to three orders of magnitude with improved accuracy. Using DISC, we reveal an inherent lack of cooperativity between cyclic nucleotide binding domains from HCN pacemaker ion channels embedded in nanophotonic zero-mode waveguides.

---

Single-molecule (SM) methods are powerful tools for providing insight into heterogenous dynamics underlying chemical and biological processes otherwise obscured in bulk-averaged measurements^1^. Use of these techniques has expanded rapidly, with modalities spanning electrophysiology, fluorescence, and force spectroscopy to probe diverse physical phenomena. Generally, SM data is obtained as a time trajectory where molecular behavior is observed as a series of transitions between a set of discrete states obscured by experimental noise. Following the growing realization that molecules involved in physiological processes exhibit a diversity of behavior, there is an increasing demand for high-throughput technologies to adequately sample various sub-populations and possible rare events^2^. Experimental advances in SM data collection such as using sCMOS cameras or nanofabricated zero-mode waveguides (ZMWs) have enabled the observation of over 1 x 10^4^ molecules simultaneously^3-5^. Non-fluorescence regimes such as plasmon rulers, magnetic tweezers, and SM-centrifugation generate a tremendous amount of data through the parallel measurement of hundreds of molecules for orders of magnitude longer recordings than fluorescence regimes, with some techniques providing week long observations of single proteins^6-9^. Despite these advances in generating statistically robust data sets, standard analysis algorithms impose a computational bottleneck at this scale of data generation^2, 3^. Here, we describe a new analysis procedure that is both accurate and orders of magnitude faster than the current methods.

Hidden Markov models (HMM) are widely used probabilistic approaches for resolving kinetics when the system states in SM trajectories are known^10-13^. Unfortunately, HMM approaches are computationally expensive and potentially inaccurate when unsupervised^14^. Model-independent approaches using unsupervised statistical learning provide an attractive alternative for uncovering dynamics without imposing constraints^15-19^. Typically, transitions are identified by change-point (CP) detection, and the total number of states is determined using hierarchical agglomerative clustering (HAC) and a specified objective function, such as Bayesian information criterion (BIC), to minimize complexity while maximizing fit (CP-HAC)^15^. CP-HAC approaches offer superior computational performance over HMM approaches when a mechanistic model is not known *a priori* but at the expense of event detection accuracy^14^. Overall, none of the existing methods can provide accurate model-free identification of states and kinetic transitions with sufficiently high computational performance to keep up with the increasing scale of data generation (**Supplemental Note 1**).

Here, we present a new algorithm for efficient and accurate idealization of SM traces in a model-independent manner. Our method, DISC (divisive segmentation and clustering), enhances CP methods by adapting divisive clustering algorithms from data mining and information theory to improve the rate and accuracy of identifying signal amplitude clusters (states) in a top-down process^20-22^. The DISC algorithm idealizes SM trajectories in three phases: 1) divisive segmentation, 2) HAC, and 3) the Viterbi algorithm (**Supplementary Fig. 1**)^23^. Together, these steps make DISC an unsupervised approach that combines the model-free state detection of CP-HAC with the high event detection accuracy of HMM at a fraction of the computational cost.

**Figure 1.**
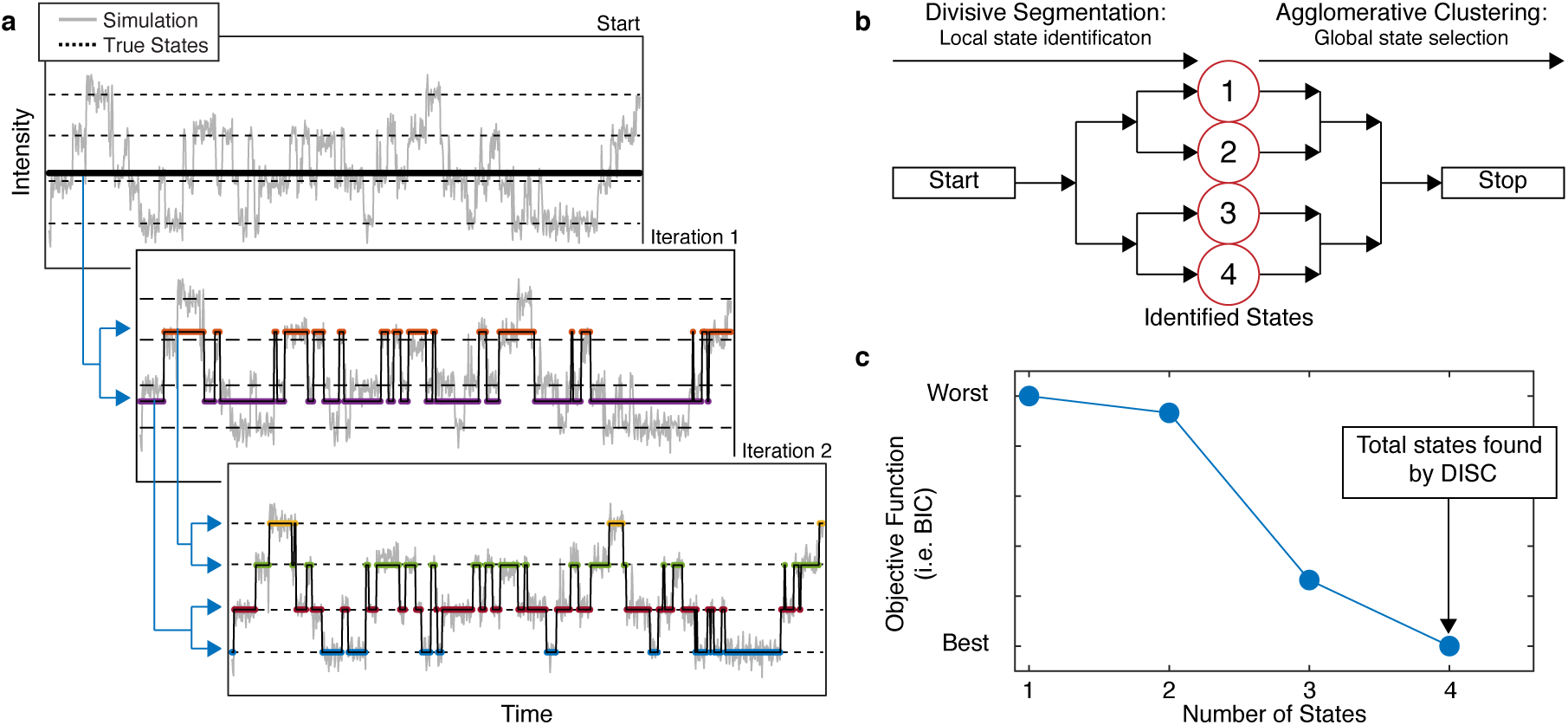
Overview of DISC. a) Stepwise discovery of change-points and states locally through divisive segmentation on a simulated trajectory featuring four simulated states (dashed lines). b) After divisive segmentation, HAC iteratively groups each identified state while computed BIC globally for the trajectory to avoid overfitting. c) The number of states with the lowest BIC value is the accepted answer for further refinement with the Viterbi algorithm.

DISC initiates with all data points belonging to one cluster and recursively partitions each cluster into two sub-clusters (**Fig 1a**). On each iteration, the cluster is denoised by applying CP analysis on the cluster’s data points (**Supplementary Note 2**). Identified intensity levels between CPs are then grouped into two sub-clusters using a modified K-means algorithm^24, 25^ The cost of increasing the complexity by splitting the original cluster into two sub-clusters is locally assessed with an objective function such as BIC which is defined in a general form for data (*D*) given a model (*M*) as:

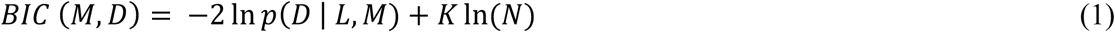

where *L* is the maximum likelihood estimate of *M, K* is the number of free-parameters in the model, and *N* is the total number of data points in the parent-cluster (**Supplementary Note 3**)^26^. If the BIC value of the parent-cluster improves with addition of two child-clusters, the split is accepted and the process repeats for the newly identified child clusters^20^. DISC continues to recursively identify sub-clusters until no cluster is further partitioned. Upon self-termination of divisive segmentation, HAC minimizes a specified objective function globally, thereby ensuring the optimum fit is obtained for the whole trace with the fewest number of states (**Fig 1b, 1c and Supplementary Fig. 2**). From these results, we estimate both the emission and transition probability matrices to identify the most probable hidden state sequence using the Viterbi algorithm, akin to HMM approaches (**Supplementary Fig. 3**)^23^.

**Figure 2.**
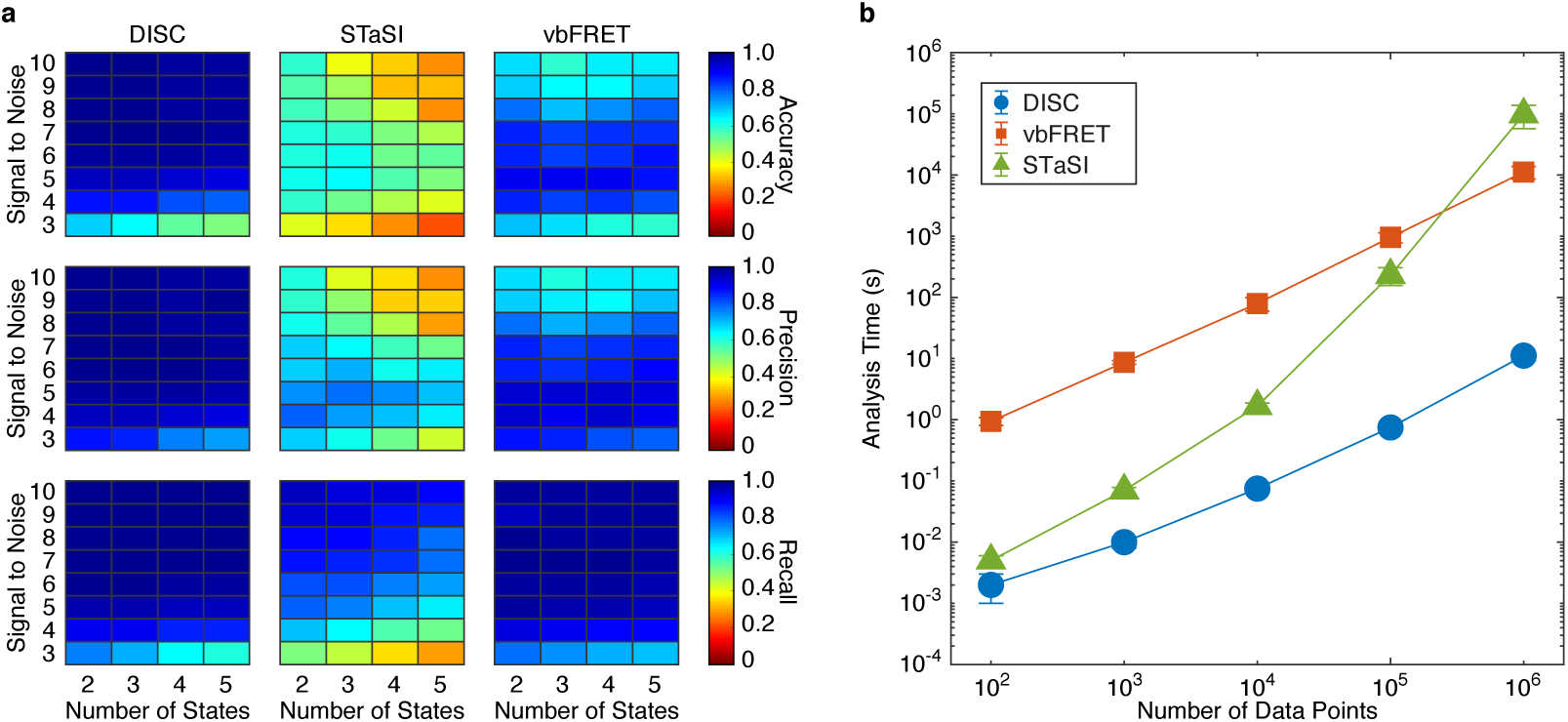
Validation of DISC on simulated single-molecule data. **a)** Average accuracy (top), precision (middle) and recall values (bottom) computed for DISC, STaSI, and vbFRET for varying SNR and number of states (**Methods**). c) Computational time (mean ± s.d.) of each algorithm for analyzing single trajectories of varying lengths. The test was performed with an Intel Xeon, 3.50 GHz processor running MATLAB 2017a.

**Figure 3.**
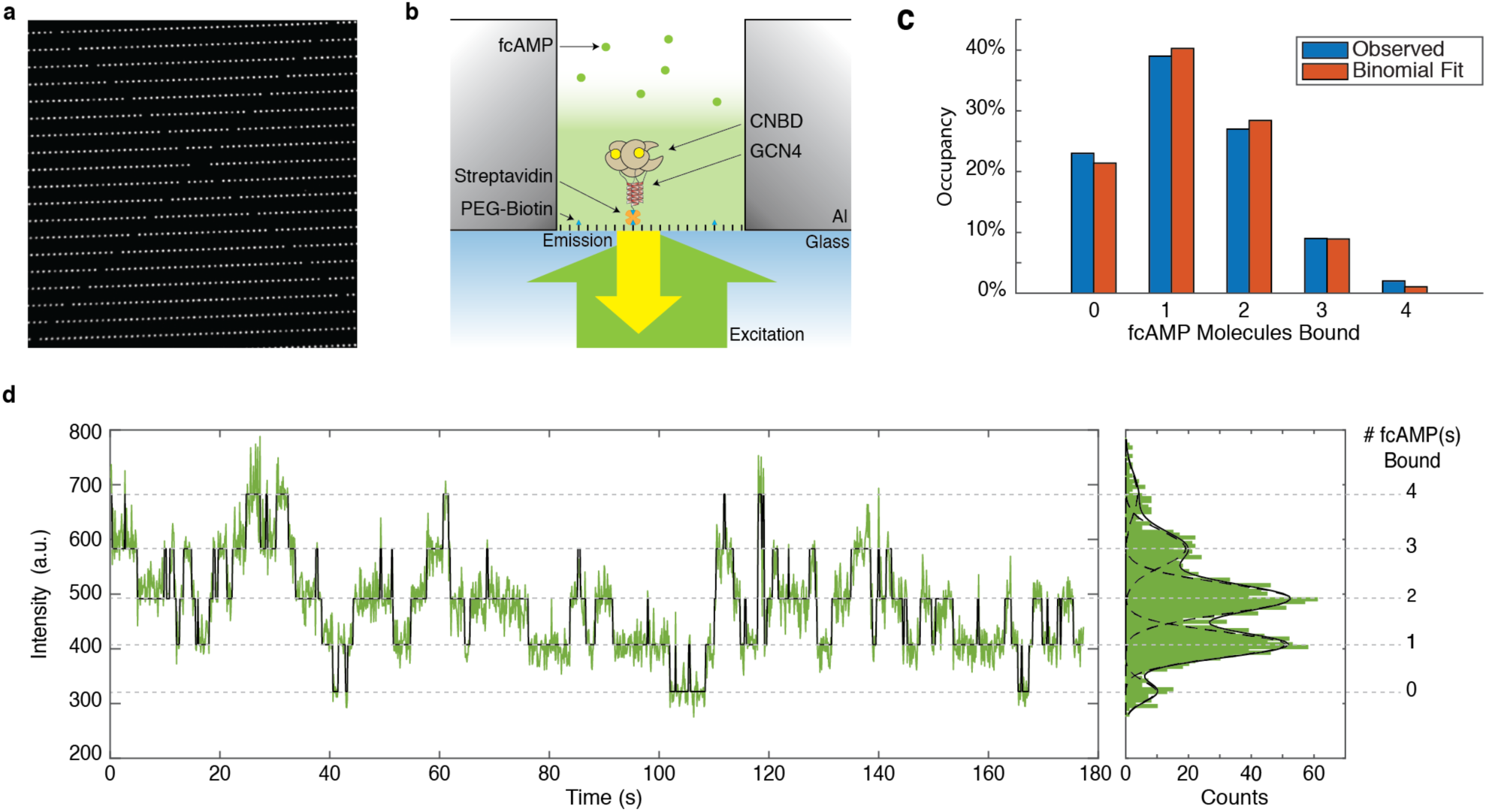
DISC reveals an inherent lack of cooperativity between HCN CNBDs. a) Representative ZMW array featuring over > 1000 ZMWs. B) Cartoon of tetrameric CNBD tethered into a ZMW for fcAMP binding experiments. c) Observed distribution of fcAMP occupancy fit with a binomial distribution. d) Representative trajectory of 1 µM fcAMP binding to tetrameric CNBD fit with DISC with up to four fcAMP molecules binding simultaneously.

We validate DISC using simulated SM fluorescence trajectories adapted from our recent studies exploring the regulatory mechanisms of cyclic nucleotide binding domains (CNBDs) from hyperpolarization-activated cyclic nucleotide gated ion channels (HCN) which regulate pacemaking in heart and brain cells (**Supplementary Note 4**)^27, 28^. By tethering isolated CNBDs into ZMWs, we can monitor binding/unbinding dynamics of fluorescent cyclic nucleotides at physiological concentrations to uncover the elementary dynamics associated with channel gating. Notably, these trajectories exhibit heterogenous bound intensity values which vary with each binding event, a common additional difficulty likely caused by shifts of the molecule in the heterogeneous excitation field or dye photodynamics (**Supplemental Fig. 5**)^5, 29^. To test the robustness of our algorithm vis-à-vis HMM and CP-HAC methods in complex kinetic landscapes, we simulated SM trajectories of ligand binding with up to four independent sites featuring heterogenous bound intensity values **(Supplemental Fig. 6)** and benchmark the results of DISC against STaSI (CP-HAC) and vbFRET (HMM) (**Methods**).

DISC maintains the highest average accuracy (0.91 ± 0.1) across 1,600 simulated trajectories (3.2 x 10^6^ data points) and is robust against false positives (precision = 0.96 ± 0.1) and false negatives (recall = 0.94 ± 0.1) at varying signal to noise ratio (SNR). (**Fig. 2a**). Although STaSI completes the analysis only 7.5X slower than DISC, it returns a low overall accuracy (0.47 ± 0.1) from frequently overfitting the number of states (precision = 0.57 ± 0.2) and missing fast transitions (recall = 0.75± 0.2) (**Supplemental Fig. 7**). While vbFRET matches the recall of DISC (0.93 ± 0.1), the analysis took over 1000X longer than DISC to complete. In addition, as the signal to noise ratio increases, vbFRET overestimates the number of states and thus lowers precision (0.8 ± 0.1) and overall accuracy (0.76 ± 0.1). Notably, DISC is the only method unaffected by inclusion of heterogenous state intensities of fcAMP (**Supplemental Fig. 8)**. When increasing trajectory duration, we find DISC is consistently 1000X faster than vbFRET and between 2X and 8700X faster than STaSI due to its quadratic time dependence^16^(**Fig. 2c**). For example, a trajectory of 10^6^ data points (common for non-fluorescence regimes) can be analyzed by DISC in 10 seconds compared to 3 hours for vbFRET and 27 hours for STaSI. Overall, DISC is both more accurate and substantially faster than current idealization approaches.

To verify performance of DISC in high-throughput collection regimes, we analyzed a large SM data set obtained with ZMWs (**Fig. 3a**). Previous studies suggest that cAMP binding to HCN channels exhibit positive-negative-positive cooperativity which may depend on membrane potential^30, 31^. To assess whether soluble CNBDs of pacemaker channels are inherently cooperative (as opposed to relying on interactions with the transmembrane domain), we examined individual binding events of fcAMP to four CNBDs linked together via a tetrameric coiled coil domain (**Fig. 3b**)^27^. Our dataset included nearly 15,000 ZMWs each monitored for 800 seconds. Each ZMW was analyzed with DISC, and the idealized trajectories were used to estimate the number of tetramers contained within a given ZMW (i.e. 0, 1, or more). By implementing cut-offs in SNR, number of identified states, and binding activity, we obtain a dataset of 287 tetrameric-CNBDs, totaling 8.76 x10^4^ seconds of combined activity, which matched our expected number of molecules (**Methods**). DISC successfully processed this entire data set in 45 minutes on a standard MacBook Air (1.6 GHz Intel Core i5) in MATLAB 2017b, whereas STaSI yielded unsatisfactory results in 5.5 hours and vbFRET would take weeks to complete (**Supplementary Fig. 9**).

To assess if the binding of cAMP to CNBDs is cooperative, we fit the fractional occupancy of each bound condition (0 to 4 fcAMPs bound) across all molecules with a binomial distribution (**Methods, Fig 3C**). Our fit returns the probability of occupancy of 1 µM fcAMP to a single CNBD to be 0.32, matching our findings with monomeric CNBDs^27^. Given that the expected binomial distribution closely resembles our experimentally obtained state-occupancy distribution, we conclude that CNBDs of pacemaker channels bind cAMP independently. This result demonstrates that the macroscopically observed cooperativity is dependent on the transmembrane domain and is not an intrinsic CNBD property.

In summary, DISC is more accurate than current widely applied idealization algorithms for analysis of SM trajectories and provides a dramatic improvement in computational speed. Consequently, DISC is an enabling technology for unsupervised processing of large data sets that allows data analysis to catch up to the enormous quantity of data available in new high-throughput SM experiments. Furthermore, DISC is modular, with modifications to CP detection and objective functions easily made to best idealize a variety of data types including fluorescence, electrophysiology, and force spectroscopy (**Supplementary Note 5**).

## METHODS

### Tetrameric CNBD preparation

The expression, purification, biotinylation, and fluorescence labeling of tetrameric CNBDs were performed as previously described^27^.

### Single-molecule fluorescence microscopy in ZMWs

Non-commercial arrays of ZMWs (Pacific Biosciences) with PEG-Biotin surface were incubated with 0.05 mg/mL streptavidin (Prospec, cat # PRO-791) for 5 minutes in a buffer containing: 40 mM HEPES, 600 mM NaCl, 20% glycerol, 2 mM TCEP, 0.1 % LDAO (Sigma, cat # 40236), 2 mg/mL bovine serum albumin (BSA), 1mM Trolox, 2.5 mM protocatechuic acid (PCA), pH 7.5 (Buffer A). After incubation, the ZMW chip was thoroughly rinsed with Buffer A to remove unbound streptavidin. Next, biotinylated tetrameric-CNBDs were diluted in Buffer A with the addition of the PCA/ PCD oxygen scavenging system by adding 250 nM of protocatechuate 3,4-dioxygenase (PCD) from Pseudoomonas sp. (Sigma, cas no. 9029-47-4)^32^ to between 100 pM and 2nM for surface immobilization in ZMWs (Buffer B). This resulted in ~100 occupied ZMWs per field of view (~1000 ZMWs) identified by fluorescence bleach steps of DY-650 that labels each of the four CNBDs. Fluorescently labeled cAMP (fcAMP; 8-(2-DY-547]-aminoethylthio) adenosine-3’,5’-cylic monophosphate) (BioLog, cat # D 109) was added at 1 µM for all single-molecule experiments in Buffer B.

ZMW arrays were placed on top of an inverted microscope (Olympus IX-71, 100X, NA 1.49) and imaged under 532 nm (60 W/ cm^2^) or 640 nm (25 W/ cm^2^) (Coherent). Notably, these experiments were conducted without FRET as previously described and therefore not limited in duration by photobleaching of the acceptor^27^. We excited DY-650 with 640 nm to identify ZMWs featuring DY-650-labeled tetrameric-CNBDs until all observable molecule photobleached. Next, fcAMP was continuously imaged with 532 nm for 8000 frames to monitor binding activity. All emission spectra were split with a 650 nm long pass dichroic (Semrock Brightline FF650) and bandpass filtered using pairs of edge filters (532-623.8 nm, 632.9-945 nm; Semrock Cy3/Cy5-A-OMF) and imaged onto two separate EMCCDs (Andor iXon Ultra X-9899) at a frame rate of 10 Hz. Excitation was controlled with Metamorph software (Molecular Devices). All data was collected using ZMWs of 150-200 nm diameter.

### Single-molecule ligand binding image analysis

All analysis was performed using custom software programs written in MATLAB (Mathworks) or ImageJ. Single-molecule trajectories of each ZMW were extracted from tiff stacks saved by Metamorph software using MATLAB. Locations of ZMWs were obtained from a threshold mask of the brightfield image of the whole ZMW array. ZMW locations were refined with a 2D Gaussian fit to the local intensity height map. The time-dependent fluorescence at each ZMW was obtained by projecting the average image intensity in a 5-pixel diameter circle onto the ZMW location throughout each image in the stack. Time series of fcAMP binding/ unbinding were analyzed with DISC.

### Trace selection and analysis of binding activity

A total of 14,937 individual ZMWs (1.2×10^8^ data points) were processed from the single-molecule tetrameric-CNBD experiments (**Supplemental Fig. 9a**). Each trajectory was idealized with DISC using a 95% confidence interval for CP detection and BIC for state selection.

sTraces featuring between 4-6 identified states with SNR ≥ 3 were retained for analysis. SNR was determined by:

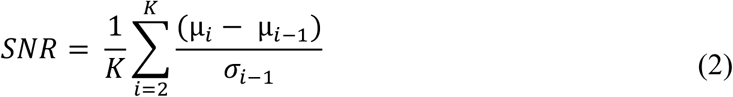

where K is the total number of states, µ is the mean intensity value of the state, and σ is the standard deviation of the data points belonging to the state. Next, we assessed the binding activity of each trace to ensure functional proteins were included in the ZMW. Using the results of monomeric CNBD studies as a guide, we selected only those traces featuring a total fraction bound (i.e. time spent in a state other than state 1) of 0.3 or greater. These simple parameters reduced the total number of viable ZMWs to 1457 compared to the 2112 total expected ZMWs identified by photobleaching steps of DY640-tetrameric-CNBDs. Critically, this reduction was not performed in order to ease the data analysis, but to confine the analysis to functional proteins. In addition, we noticed an asynchronous decay of protein activity over excitation time (**Supplemental Fig. 10).** We are unsure as to whether this is due to the protein entering a non-binding conformational state or detaching from the surface. To account for this, we binned the activity of each trajectory into 1000 frames (of 8000 frames total) and calculated the fraction bound per bin. Traces that exhibited long time periods without binding activity (< 0.3 average fraction bound) were truncated within the bin using the CP method to identify the most likely frame where protein behavior changed. Thus, only the frames wherein protein appears receptive to fcAMP were analyzed. Traces were refit by DISC to account for variance in truncation and re-selected by SNR and number of states. This procedure left 420 ZMWs for visual inspection, from which a total of 287 tetrameric-CNBDs exhibiting four or five conformational states (3 to 4 fcAMPs bound) were included for analysis. In total, 8.76×10^4^ seconds of combined activity featuring 3.77 × 10^4^ events (**Supplementary Fig. 11**) was analyzed. Binomial fitting of the fractional occupancies spent in each state was performed using MATLAB’s mle function.

### Single-molecule simulations

Single-molecule trajectories were simulated as a Markov process of transitions between discrete states. All simulations featured frame rate of 10 Hz and variable durations, SNR, and number of states. The kinetic scheme used was adapted from our recent studies of fcAMP binding to isolated monomeric CNBDs^27^. This model is a four-state scheme where both the unbound (U) and bound states (B) exhibit conformational changes (U’ ⇔ U ⇔ B ⇔ B’), yet exhibit only two different emissive states. fcAMP binding occurs between U and B. The rate constants (s^-1^ or M^-1^ s^-1^) are: k_U’U_ = 0.15; k_U,U’_ = 0.04; k_U,B_ = 0.23 * [fcAMP]; k_B,U_ = 0.95; k_B,B’_ = 0.51; k_B’,B_ = 0.31 at 1 µM fcAMP. To mimic the tetrameric nature of HCN channels with no cooperativity, we extrapolated up to four bound states by summing independent CNBD trajectories prior to the addition of noise. State intensities values and heterogenous bound intensity emissions were each drawn from lognormal distributions fit to monomeric CNBD single-molecule data, with average intensities between subsequent states being uniform. (**Supplemental Figs. 4-6**). Gaussian noise was applied at trajectories to the specified SNR.

DISC, STaSI, and vbFRET are all written entirely in MATLAB (MathWorks). Each algorithm was used outside of their graphical user interfaces (GUIs) to more accurately compare the computational time of native functions within each algorithm. User parameters in DISC include: the confidence interval of CP detection and the objective function(s) for clustering. Unless otherwise stated, a 95% confidence interval was applied for CP detection and BIC was used for all clustering. Changing these values can result in different results and will be dependent on the type of data being analyzed (**Supplemental Note 4**). For analysis with STaSI and vbFRET, we used the recommended default values set by their authors.^11, 16^ For STaSI, this means a 99.8% confidence interval of CP detection. In vbFRET, users must provide the number of states and fitting attempts per trace (left at the default value of 10). In order to use vbFRET in an unsupervised manner, we wrote a wrapper around the provided *vbFRET_no_gui.m* script to perform analysis outside of the vbFRET GUI. The wrapper begins by fitting the trace to one state and increases the number of states until two more beyond the number of states with the maximum evidence to ensure the maximum fit has been obtained. As no changes were made to native vbFRET functions, implementing this wrapper has no effect on vbFRET’s accuracy. We expect changing parameters in both STaSI and vbFRET may lead to different results; however, it was not our goal to optimize the use of these algorithms.

All quantifications of computational time were performed using the tic and toc functions in MATLAB. For idealization accuracy, each event returned by a given algorithm is classified as a True Positive (TP), False positive (FP), or False Negative (FN). We define a TP as being in the correct state (± 10 % the correct intensity level) and correct event duration (± 1 frame) for a given simulated event. FPs are either added events or correct events in the wrong state and FNs are missed events. For each trajectory, we computed accuracy, precision, and recall as:

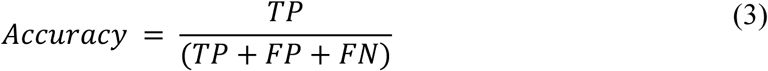

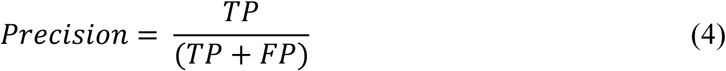

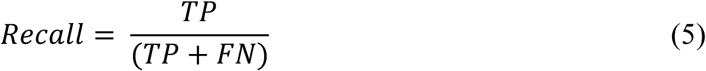

Accuracy represents the overall performance, whereas precision and recall highlight the false positive error rate (overfitting the data) and false negative rate (underfitting the data), respectively.

## Acknowledgments

We thank Dr. Mike Sanguinetti for the wild-type HCN2 plasmid and Dr. Vadim A. Klenchin for the purification of the tetrameric-CNBD. We also thank Owen Rafferty for his assistance in the development of the GUI for running the DISC algorithm. This research was supported by the NIH grants to B.C (NS-101723, NS-081320, and NS-081293), D.S.W (T32 fellowship GM007507) and R.H.G. (GM127957).

## Contributions

D.S.W. conceptualized and wrote the DISC algorithm. D.S.W and M.P.G.O performed and analyzed SM experiments. D.S.W and M.P.G.O wrote all MALTAB scripts for the image processing, SM simulations, and algorithm quantifications. D.S.W., M.P.G.O., R.H.G, and B.C. contributed to conception, experimental design, and writing of the manuscript.

## Code Availability

DISC is written in MATLAB both with and without a user-friendly GUI. The DISC package, along with scripts for simulating multi-ligand binding data and quantification of idealization results will be made available once published on https://github.com/ChandaLab.

## Data Availability

Raw and simulated SM data that support these findings are available from the corresponding authors upon request.

