## Supplemental Information for "High-Throughput Single-Molecule Analysis via Divisive Segmentation and Clustering"

**Supplementary Table 1. Common Abbreviations**

| Abbreviation | Description |
|---|---|
| BIC | Bayesian Information Criterion |
| CI | Confidence Interval (of change-point detection) |
| CNBD | Cyclic nucleotide binding domain |
| CP | Change-point |
| CP-HAC | Use of change-point detection and hierarchical agglomerative clustering to minimize an objective function for state determination |
| DISC | Divisive Segmentation and Clustering |
| EMCCD | Electron multiplying charge-coupled device |
| fcAMP | Fluorescent cyclic adenosine monophosphate |
| FRET | Förster resonance energy transfer |
| GUI | Graphical User Interface |
| HAC | Hierarchical agglomerative clustering |
| IC | Information criteria (used interchangeably with "objective function") |
| MDL | Minimum Description Length |
| HMM | Hidden Markov model |
| smFRET | Single-molecule FRET |
| SM | Single-Molecule |
| SNR | Signal to noise ratio |
| STaSI | State transition and step identification |
| vbFRET | Variational Bayes FRET |
| ZMW | Zero-mode waveguide |

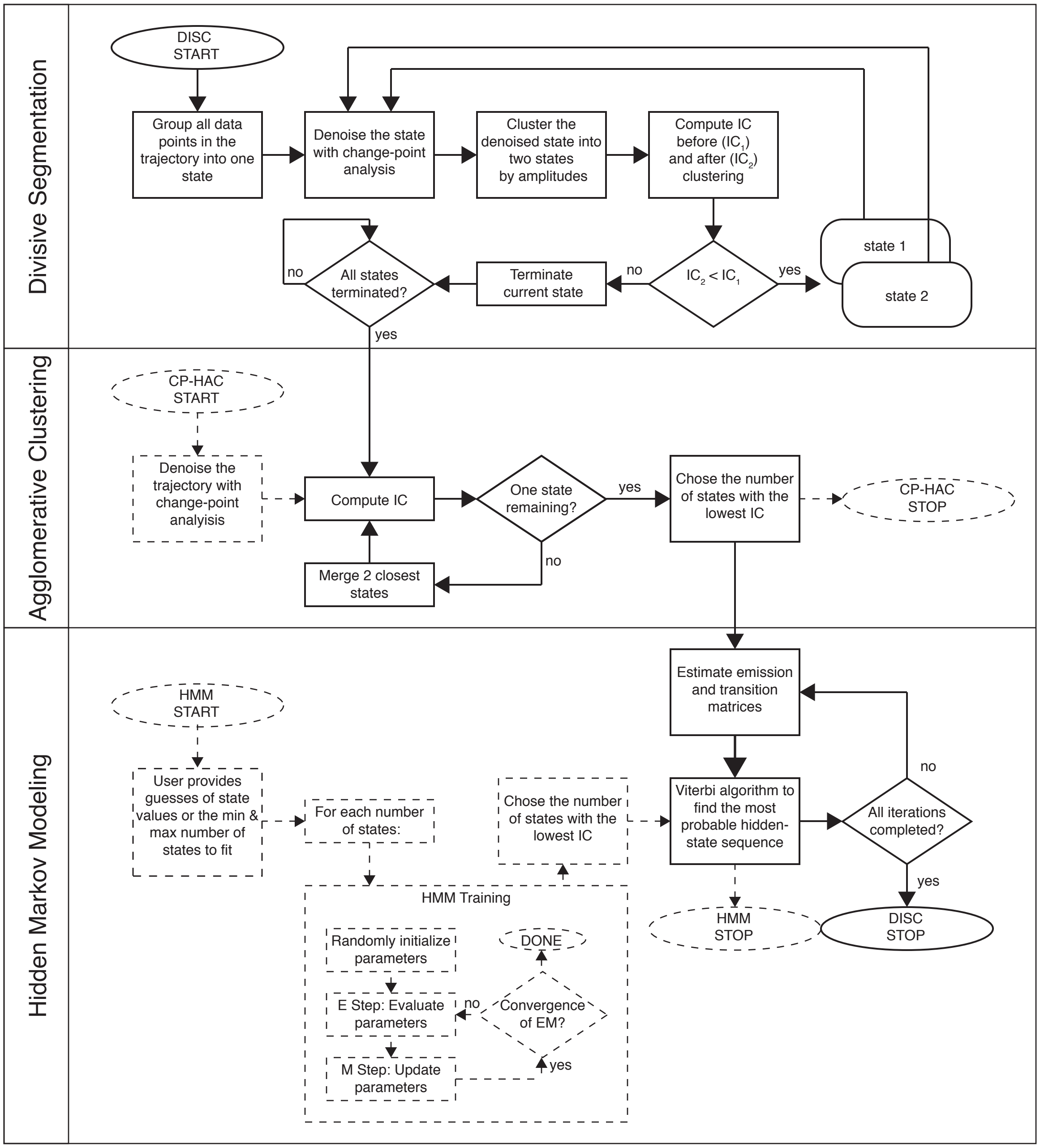


**Supplementary Figure 1. Workflow of DISC.** Solid lines indicate the path DISC takes through divisive segmentation, hierarchical agglomerative clustering, and hidden Markov modeling (Viterbi algorithm). IC = information criterion. General CP-HAC and HMM approaches are shown for comparison. Dotted lines indicate steps that do not overlap with DISC. Ovals indicate start/ stop; rectangles indicate a process or computation; diamonds represent decisions. Note, most current uses of HMMs involve a single user-defined number of states.

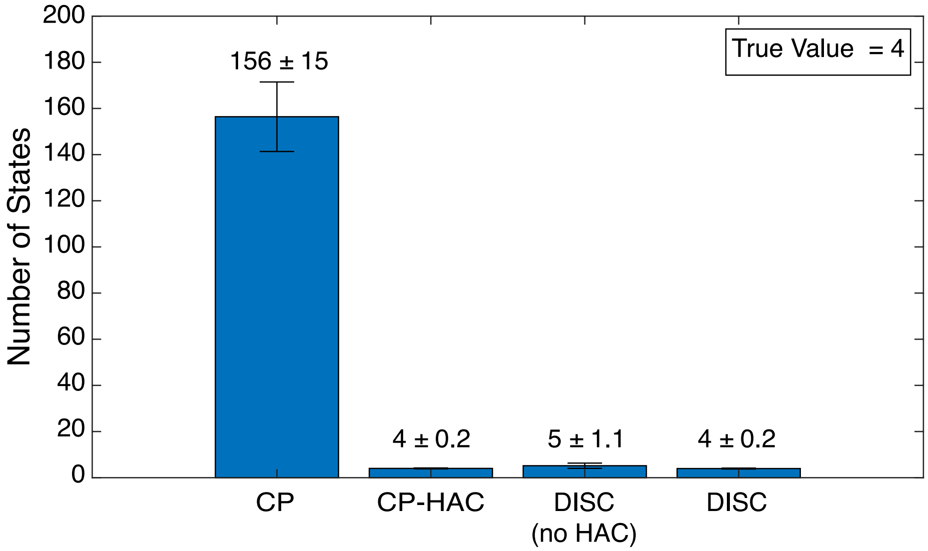


**Supplementary Figure 2: Number of states identifed by change-point detection, DISC without HAC, and DISC.** Average number of states (mean ± s.d.) across 500 simulated trajectories of 2000 frames each with a SNR = 4 and 4 total states. Values were obtained with 95% CI of CP detection and BIC for clustering. While CP-HAC and DISC both find the correct answer, DISC was 35X faster.

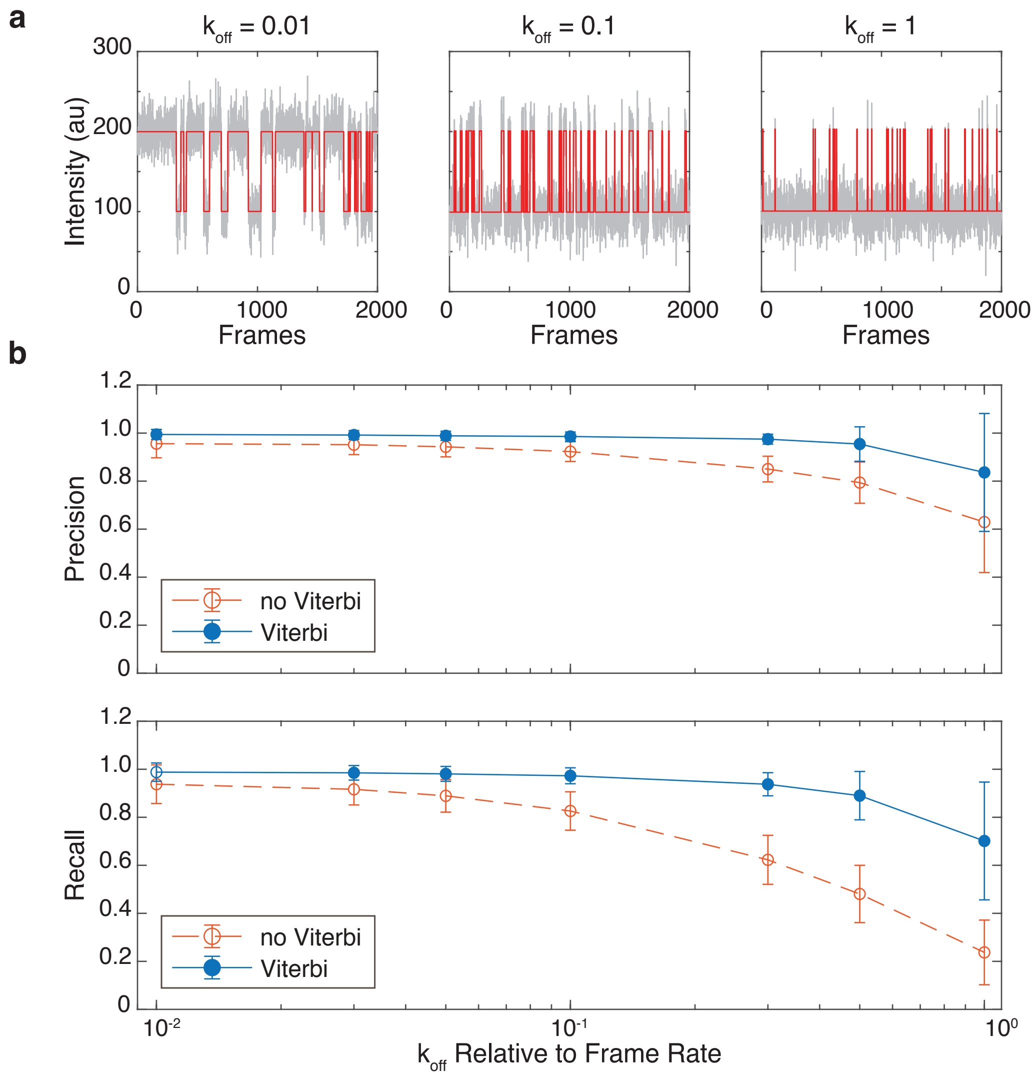


**Supplementary Figure 3: Event detection accuracy of DISC with and without Viterbi step.**

a) Examples of simulations of a two-state system with a $k_{on}$ = 0.02 frames$^{-1}$ and varying $k_{off}$. b) Values (mean ± s.d.) obtained at 95% CI of CP detection across 500 trajectories each with 2000 frames per condition. See methods for definitions.

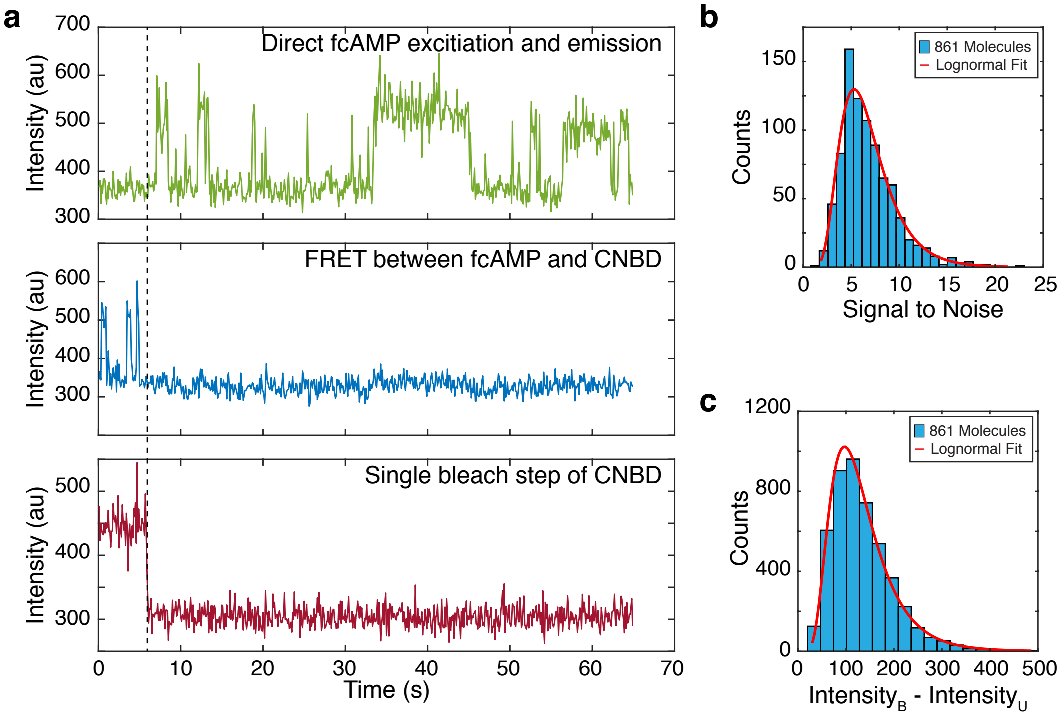


**Supplementary Figure 4. Characterization of fcAMP binding to monomeric CNBDs in ZMWs.** a) Representative experimental trajectory of a fcAMP binding to an isolated CNBD in a ZMW. (b) Distribution of signal to noise ratios of each per trajectory (N=861) with a log-normal fit of $\mu = 1.84$, $\sigma = 0.41$. (c) Distribution of the difference in intensities between bound (B) and unbound (U) states per trajectory (N = 861) with a log-normal fit of $\mu = 4.79$, $\sigma = 0.47$. Log-normal fitting was performed using MATLAB's `lognfit` function.

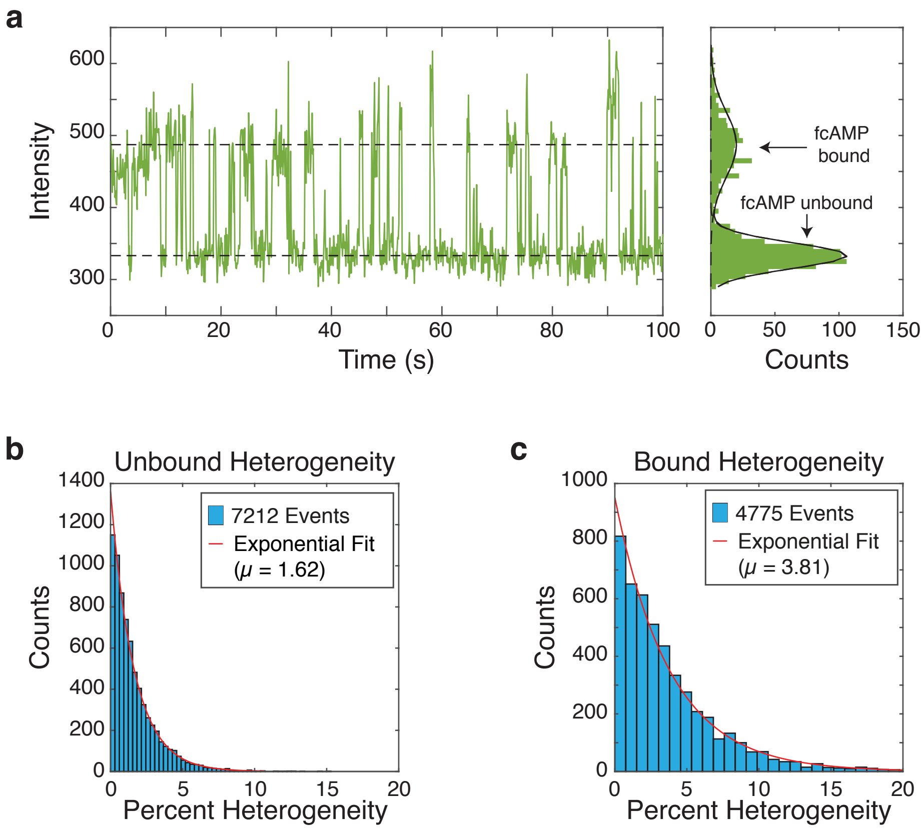


**Supplementary Figure 5. Heterogenous Intensities upon fcAMP binding.** a) Representative experimental time series of a fcAMP molecule binding to an isolated CNBD tethered in a ZMW. Across the trajectory, each individual binding event exhibits varying intensities, despite the clear appearance of two Gaussian distributions. Quantification of unbound event heterogeneity (b) and bound event heterogeneity (c) (see **Supplementary Note 3**). Log-normal fitting was performed using MATLAB's `lognfit` function.

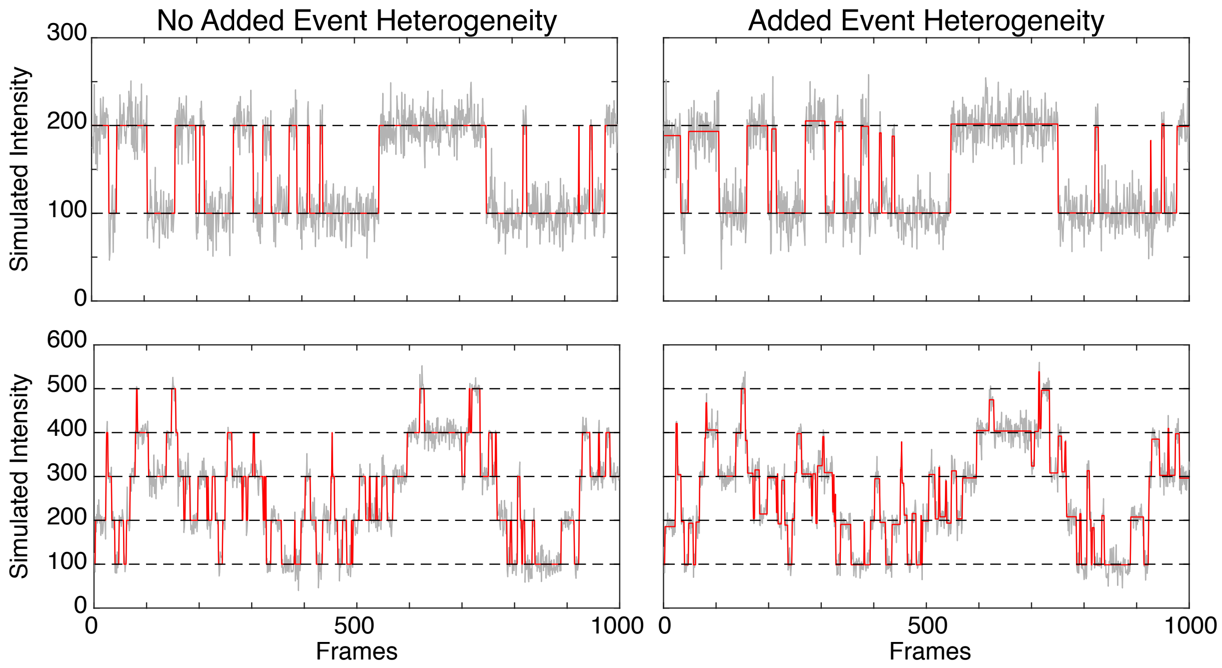


**Supplementary Figure 6. Simulations with heterogenous fcAMP emission.** Representative SM simulations without (left) and without (right) heterogenous emission of fcAMP upon binding for two states (top) and five states (bottom). Plots show simulated trajectory (red), overlaid with the addition of Gaussian noise (grey) and the average intensity value of each state (dashed black).

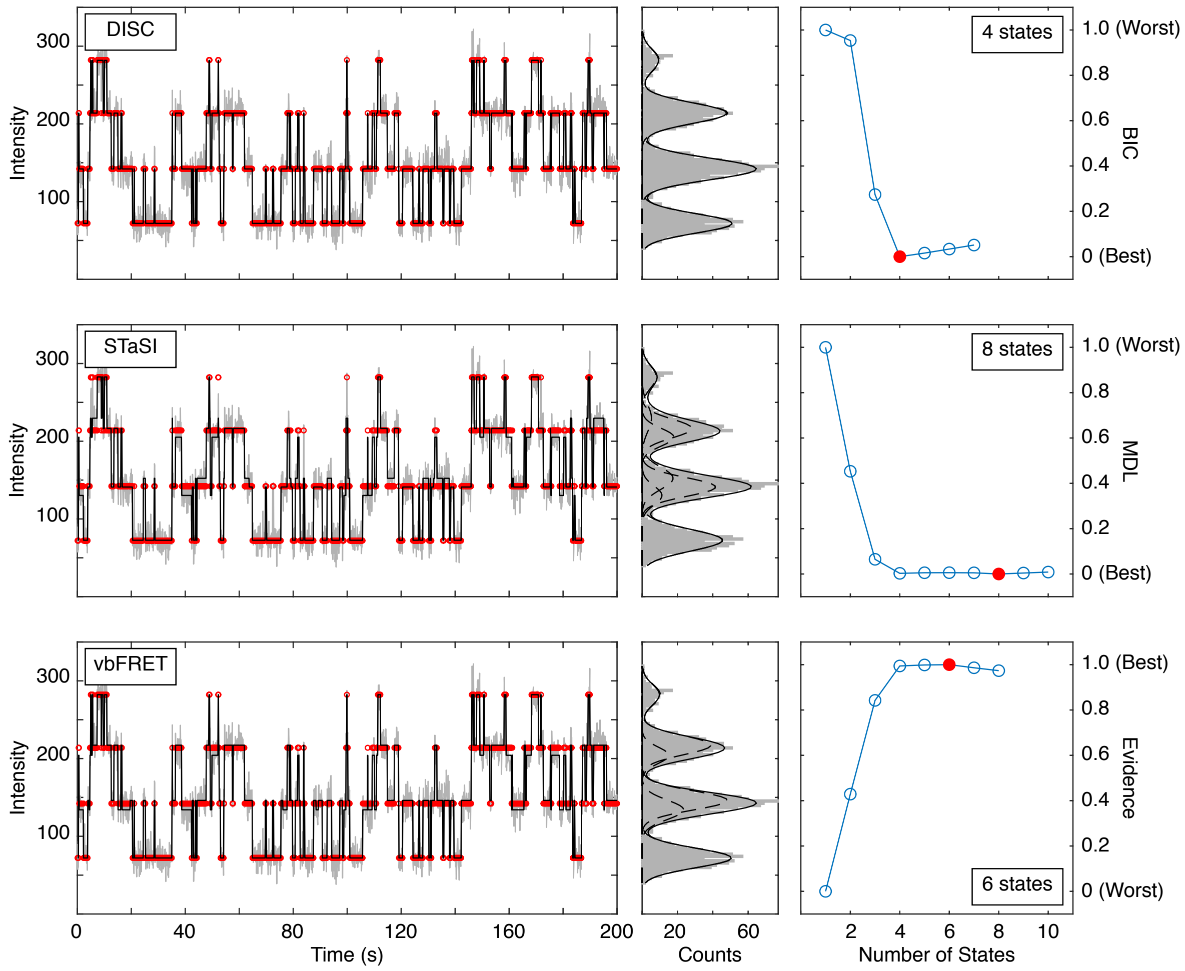


**Supplementary Figure 7. Representative fit of each algorithm on simulated data.** Simulated trajectory (red) with 4 true states fit and added Gaussian noise (grey) to SNR = 6 overlaid with fits (black) from DISC (top), STaSI (middle), or vbFRET (bottom).

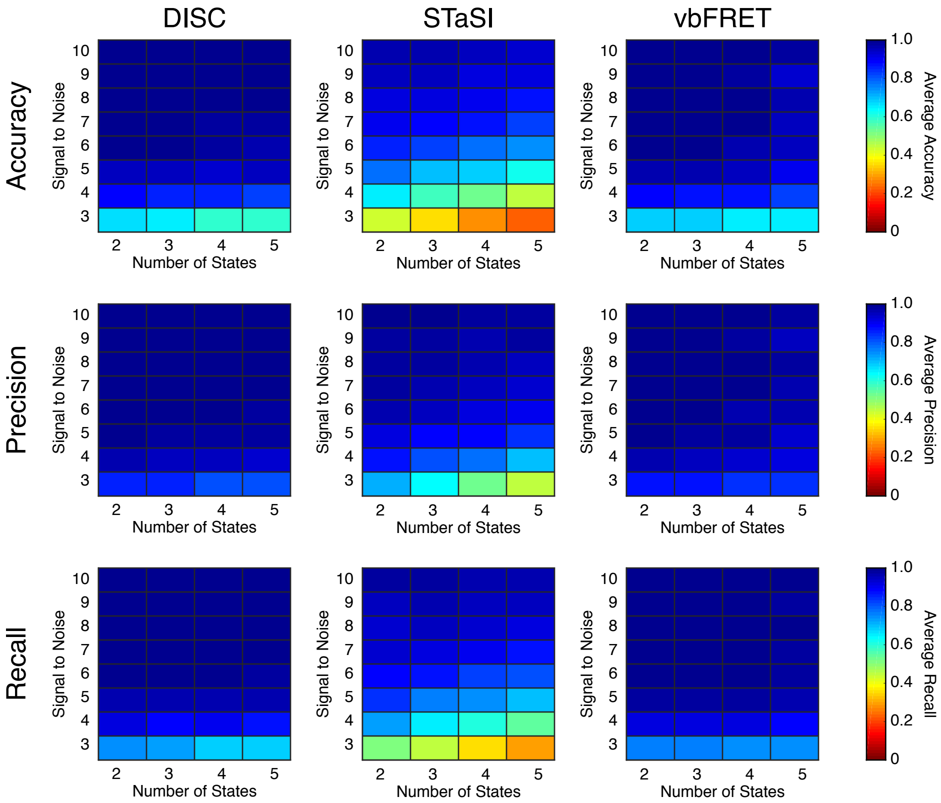


**Supplementary Figure 8. Algorithm performance on simulations without heterogenous fcAMP emission.** For each condition, 50 trajectories were simulated following for the provided number of states and SNR with 2000 frames at a frame rate of 10 Hz. In total, 1600 trajectories and 3.20 x $10^6$ data points were assessed. Heterogenous intensities per binding event were not included. Each value is the average accuracy (top), precision (middle), and recall (bottom) (**Methods**).

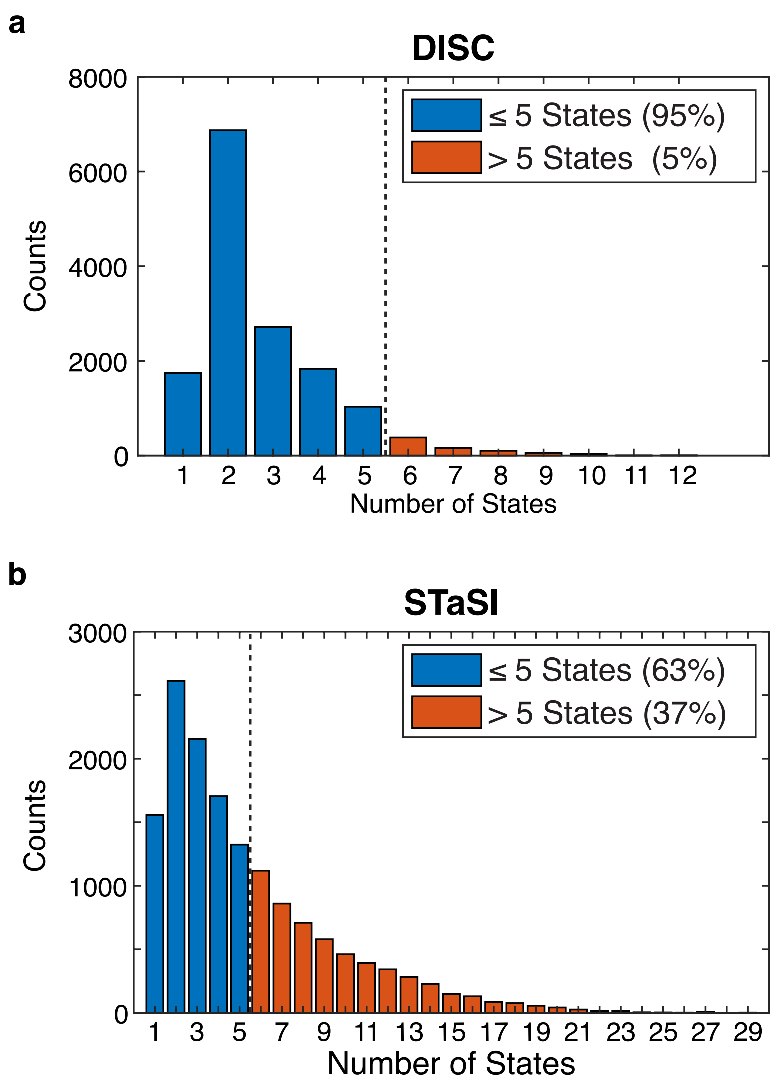


**Supplementary Figure 9. Tetrameric CNBD analysis by DISC and STaSI.** Initial number of states found by DISC (a) and STaSI (b) when run on the tetrameric CNBD data set across 14,937 trajectories prior to trace selection (see **Methods**). The expected distribution is between 1 and 5 states per trajectory to account for empty ZMWs and fully occupied tetrameric CNBDs (4 fcAMP bound states plus 1 unbound states). This analysis was not repeated using vbFRET as we estimated the process would take weeks to complete.

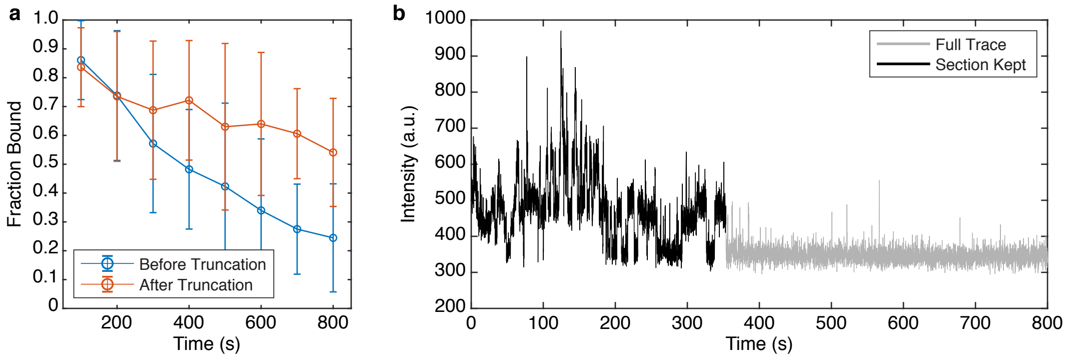


**Supplementary Figure 10. Asynchronous decay of tetrameric CNBD activity over excitation time.** a) Average fraction bound of each molecule over time (binned every 100 s) before and after trajectory truncation (**Methods**). b) Representative truncated trajectory.

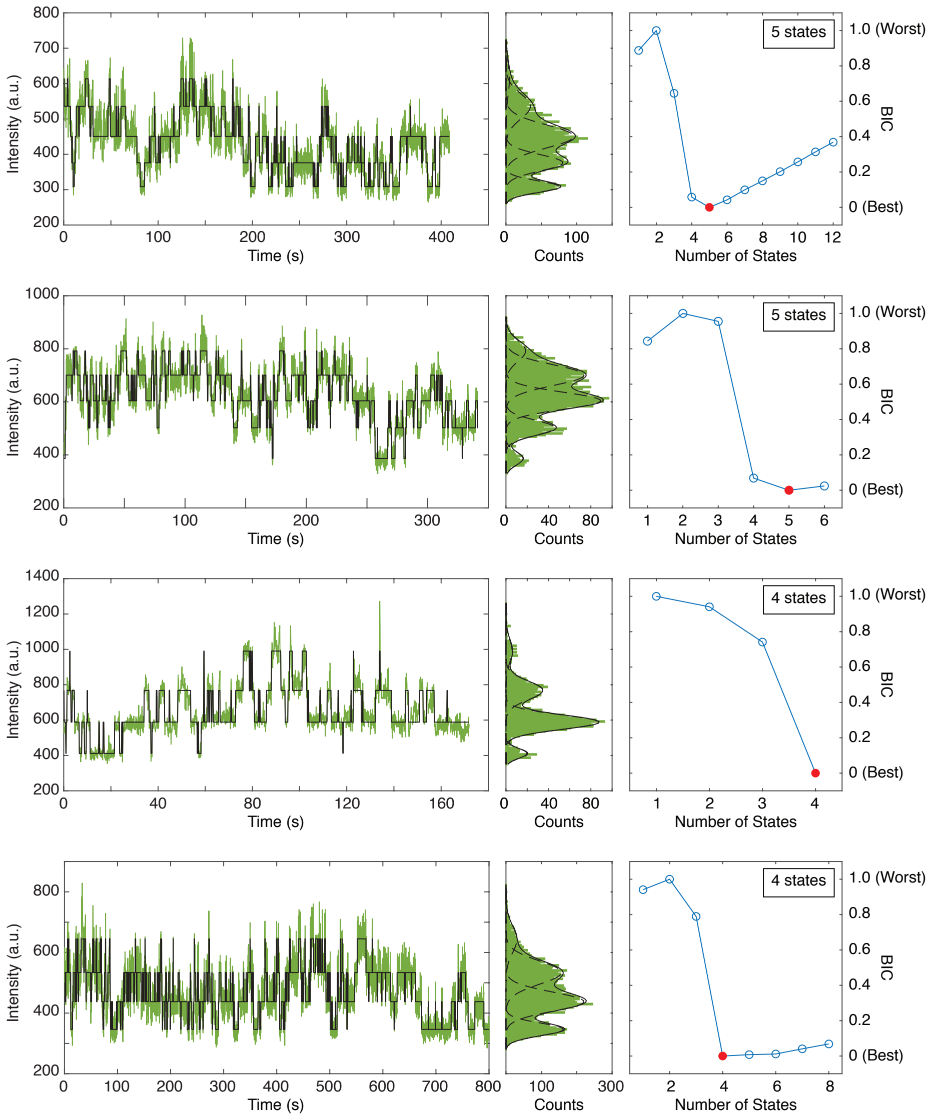


**Supplementary Figure 11. Example trajectories of 1μM fcAMP binding to tetrameric CNBDs in ZMWs.** Representative trajectories featuring up to 3 or 4 bound fcAMP molecules analyzed by DISC (left) with distribution fits (middle) and BIC curves for optimal state selection (right).

**Supplementary Note 1.** Standard SM Analysis Methods

Statistical modeling of single-molecule trajectories often adopts one of two types of approaches. The first is a probabilistic approach that models a molecule's behavior as a Markov chain, wherein the protein transitions between discrete states whose outputs are what is measured experimentally (HMM). This involves estimating the transition probabilities between a small set of postulated states with defined outputs using methods to maximize the likelihood of the model given the observations or Bayesian inference to estimate model parameter distributions. The use of HMMs for SM analysis was pioneered in the ion channel community through software such as QuB[1,2], which provides a flexible GUI for users to build models and analyze trajectories. Notable, QuB implements the segmented K-means algorithm, which is a fast method for estimating the parameters of the user provided model[3]. The first HMM software designed especially for smFRET analysis was HaMMy and has been widely used since its 2006 release[4]. The adaptation of variational Bayesian inference for HMM training of smFRET data was deployed in vbFRET[5], which provides a simple GUI for the analysis of smFRET trajectories at much faster speeds than QuB or HaMMy[6].

Although powerful, HMMs are most commonly used in a supervised regime where the user postulates model parameters such as the number of states, their measured outputs, and the allowed transitions between them. As this information is often not known *a priori*, it is desirable to test multiple models and rank them according to Bayesian probabilistic approaches or objective functions such as BIC. This process can dramatically increase the analysis time to ensure the parameters space has been fully explored. Improvements to this process are still under active research development, with approaches such empirical Bayesian FRET (ebFRET)[7] and infinite

HMMs using Bayesian nonparametric inference[8,9] to naturally learn the states in the trajectory without the typical parametric model selection.

The second type of approach includes model-independent CP detection methods which often rely on the popular binary separation approach[10]. On each iteration, a single CP (i.e. temporal point of a transition between states) can be detected by hypothesis testing each data point to assess whether or not a transition has occurred. Each identified CP divides the time series into two segments and the process is recursively repeated on each segment until no further CPs are detected given the provided confidence interval for the analysis. Upon completion, segments between identified CPs are iteratively grouped in a bottom-up fashion using a greedy hierarchical agglomerative clustering (HAC) algorithm until one segment remains. Thereafter, an objective function is used to select the best number of states for the trajectory. The pioneering application of CP-HAC to SM data entailed sequential use of a Poisson merit function for CP detection and expectation-maximization clustering which enabled the determination of states at superior accuracy over the common threshold and binning approaches used at the time[11]. Variants to this framework include approaches such as STaSI which adapted the use of the Student's T-test for rapid CP detection and MDL for state selection[12].

A major advantage of the CP-HAC methods is that they only require a confidence interval and/or an objective function for the analysis, unlike HMMs which require a model to fit. Further, they have broad applications in non-linear piecewise data such as processive SM data[13]. Unfortunately, CP-HAC methods exhibit a quadratic time complexity[12]. While computational tricks have been employed to speed up the process, such as the use of parallel processing[11], CP-HAC methods are typically used for the shorter trajectories common in SM fluorescence experiments, such as smFRET, and would be difficult to apply for longer time traces such as those found in single-channel recordings or in fluorescence microscopy with fluorogenic reactions. For a concise introduction of the application of unsupervised learning for SM analysis, we refer readers to Li and Yang (2019)[14].

**Supplementary Note 2.** Change-Point Detection in DISC

Change-points are identified using a recursive binary segmentation algorithm. For N data points contained in vector of data X [$X_1$, $X_2$…, $X_N$], a hypothesis test is conducted at every data point *k* to determine if a changepoint has occurred between $X_1$:$X_k$ and $X_{k+1}$:$X_N$. To assess if there is a significant difference in the mean intensities between these two segments, we use a Student's t-test of unequal sample size but uniform variance akin to STaSI and compute t by:

$$t = \frac{|\mu_1 - \mu_2|}{\sigma\sqrt{\frac{1}{x} + \frac{1}{N-x}}} \tag{1}$$

where $\mu_1$ is the average intensity of the data points from 1 to *k*, $\mu_2$ is the average intensity of all data points from *k* + 1 to N, and σ is the standard deviation of noise which is estimated using a Haar wavelet transform (see the Supporting Information of Shuang et al., for a detailed derivation)[12]. The largest t-value computed for the set of data points is then compared to the critical value corresponding to the confidence interval provided by the user (for example, a 95% confidence interval has a critical value = 1.96). If t > critical value, the change point is accepted and the data is segmented at the change-point. This process recursively repeats to identify change points in each new segment until no significant change-points are found.

**Supplementary Note 3.** Hierarchical Agglomerative Clustering and State Determination

After divisive segmentation, all identified states are iteratively grouped using a greedy HAC algorithm until one state remains. Here, we determine the similarity between neighboring clusters A and B by Ward's distance[15], which considers the number of data points in each state (n) and the Euclidean distance between the means of the states by

$$d(A,B) = \sqrt{\frac{2n_A n_B}{n_A + n_B}} \|\mu_A - \mu_B\|_2 \tag{2}$$

Notably, while divisive segmentation tends to slightly overfit the number of states, CP detection alone can often identify orders of magnitude more unique states (**Supplementary Figure 2**). Overall, we find sequential use of divisive segmentation and HAC to be a more accurate scheme for state identification.

To determine the optimal number of states, an information criterion (or objective function) is computed on each iteration of HAC to maximize the fit of the trajectory while minimizing complexity. As discussed in the main text, we implement the BIC, which is defined in a general form as:

$$BIC\ (M,D) = -2\ln p(D \mid L, M) + K \ln N \tag{3}$$

where M is the model, D is the data, L is the likelihood of M, K is the number of parameters in M, and N is the number of data points. Following Bishop, in the case of a single data point x, we can evaluate a normal distribution by:

$$\mathcal{N}(x \mid \mu, \sigma) = \frac{1}{\sigma\sqrt{2\pi}} e^{\frac{-(x-\mu)^2}{\sigma^2}} \tag{4}$$

In the cause of a mixture of 1D Gaussians, we can evaluate each as a linear combination of each

K Gaussian density[16]. Therefore, the p(x|M) is the sum of all probabilities across K components, where each component (j) is characterized by a mean ($\mu_j$), standard deviation ($\sigma_j$) and mixing coefficient ($\omega_j$) .

$$p(x \mid M) = \sum_{j=1}^{K} \omega_j * \mathcal{N}(x \mid \mu_j, \sigma_j) \tag{5}$$

For the BIC calculation, M is described by K components, each featuring values ($\mu$, $\sigma$, $\omega$). Given that $\sum_{j=1}^{K} \omega_j = 1$, the overall degrees of freedom for fitting the model is 3K–1. Therefore, for N data points contained in vector X [$X_1$, $X_2$…, $X_N$],

$$BIC = -2 * \ln\left[\sum_{j=1}^{K} \omega_j * \frac{1}{\sigma_j\sqrt{2\pi}} e^{\sum_{i=1}^{N}\left[-(X_i - \mu_j)^2/\sigma_j^2\right]}\right] + (3K - 1) * \ln(N) \tag{6}$$

**Supplementary Note 4.** Analysis of Monomeric CNBDs in ZMWs

We previously explored the binding mechanism of fcAMP isolated CNBDs labeled with DY-650 in ZMWs[17]. In these experiments, we simultaneously monitored: 1) the DY-650 CNBD photobleaching step, 2) FRET between the donor fcAMP molecule and the CNBD (ZMW-FRET), and 3) the direct occupancy of the fcAMP molecule (ie, a one-color fluorescence measurement). While the use of ZMW-FRET can resolve binding dynamics at up to millimolar concentrations of fluorescently labeled molecules[18], the latter approach enables substantially longer recording times unshackled from the limitation of acceptor photobleaching, a scheme that was necessary to resolve the binding of all four fcAMP molecules to tetrameric-CNBDs. Therefore, we quantified the direct fcAMP excitation/ emission trajectories following acceptor photobleaching from the monomeric

CNBD data set used in our previous work to use a basis for our tetrameric-CNBD studies (**Supplementary Figure 4**)[17]. The data set consisted of 861 single molecules for a combined acquisition time of 44,090 seconds (4775 total binding events). All trajectories had a SNR > 2.

To quantitate the heterogenous intensities from fcAMP binding (**Supplementary Figure 5**), the mean of individual bound event intensities was taken for each identified event, so long as the event was > 2 frames in duration. Heterogeneity was computed as the absolute percent different of each event vs the mean bond intensity for the given trajectory by:

$$Percent\ Heterogenity = \left| \frac{< I >_{event} - < I >_{Bound}}{< I >_{Bound}} \right| \times 100\% \qquad (9)$$

The obtained distributions of SNRs and heterogenous event intensities were used for simulations of tetrameric-CNBDs in the main text (**Supplementary Figure 6**). The heterogeneity of unbound events was also assessed (**Supplementary Figure 6b**) but was deemed very subtle to and was therefore not included in any simulations.

**Supplementary Note 5.** Changing Parameters in DISC

As briefly discussed in the main text, we find the use of BIC derived from a mixture of Gaussians performs well in finding the optimal number of states in simulated binding data. We do not claim that the parameters used for this analysis (95% CI for CP detection and BIC for state selection) will be optimal for all data-sets across all experimental modalities. For example, slower dynamics will likely be overfit by a low CI while fast dynamics will be missed by a high CI. We have intentionally developed DISC as a flexible framework such that different hypothesis tests of CP detection and clustering may be implemented depending on the type of data being used. While we explored the use of the Student's T-Test for change-point detection for EMCCD noise, the use of merit functions for Poisson or Gaussian distributions may be more applicable to different data sets[11, 14, 19]. The same holds true for the use of a Gaussian BIC for cluster determination, as other information criterions and objective functions will likely perform with different results. For example, we speculate that for primarily Poisson data, such as single ion-channel data or data obtained from an Avalanche photodiode (APD), the use of a Poisson merit function for CP detection and a Poisson derived BIC would be more applicable[19]. In addition, we speculate that state identification in SM data of processive trajectories would be better suited using objective functions such as MDL[12], which minimize residuals across the trajectory rather than distribution fitting approach of BIC which would likely underfit the number of states. Such modular changes can be easily implemented in the DISC framework.
